# A methodology combining reinforcement learning and simulation to optimize the *in silico* culture of epithelial sheets

**DOI:** 10.1101/2023.04.25.538212

**Authors:** Alberto Castrignanò, Roberta Bardini, Alessandro Savino, Stefano Di Carlo

## Abstract

Tissue Engineering (TE) and Regenerative Medicine (RM) aim to replicate and replace tissues for curing disease. However, full tissue integration and homeostasis are still far from reach. Biofabrication is an emerging field that identifies the processes required for generating biologically functional products with the desired structural organization and functionality and can potentially revolutionize the regenerative medicine domain, which aims to use patients’ cells to restore the structure and function of damaged tissues and organs. However, biofabrication still has limitations in the quality of processes and products. Biofabrication processes are often improved empirically, but this is slow, costly, and provides partial results. Computational approaches can tap into biofabrication underused potential, supporting analysis, modeling, design, and optimization of biofabrication processes, speeding up their improvement towards a higher quality of products and subsequent higher clinical relevance. This work proposes a reinforcement learning-based computational design space exploration methodology to generate optimal in-silico protocols for the simulated fabrication of epithelial sheets. The optimization strategy relies on a Deep Reinforcement Learning (DRL) algorithm, the Advantage-Actor Critic, which relies on a neural network model for learning. In contrast, simulations rely on the PalaCell2D simulation framework. Validation demonstrates the proposed approach on two protocol generation targets: maximizing the final number of obtained cells and optimizing the spatial organization of the cell aggregate.

## 1. Introduction

Biofabrication, involving the automated creation of biologically functional products from living cells and biomaterials, aims to replicate the complexity of biological systems, from cell aggregates to organs [1, 2]. Tissue Engineering (TE) and Regenerative Medicine (RM) leverage patient cells to repair damaged tissues and organs [1], yet achieving full tissue integration remains challenging [3]. The design of biofabrication processes is complex, with many variables influencing product quality [4]. Traditional Design Space Exploration (DSE) is costly and often relies on human effort or chance, despite advancements in automation and digitalization [5].

Established experimental designs like One Factor At a Time (OFAT), factorial design, and Design of Experiment (DoE) help optimize resource use in biological experiments [6, 7, 8], but DoE struggles with the vast experiments needed in complex fields like biofabrication [9, 10, 11].

Computational and Artificial Intelligence (AI) methods offer advanced support in biofabrication design and optimization [12, 13]. They enable systematic DSE, considering complex parameter interactions and process outputs. Effective computational approaches require white-box models that represent biological processes accurately [14]. However, current models often neglect biological complexity, leading to large state spaces and necessitating metaheuristics for feasibility [15, 16].

Previous research applied a model-based Optimization via Simulation (OvS) approach using Genetic Algorithm (GA) for the biofabrication of epithelial sheets [17, 18]. This paper extends this exploration, applying Machine Learning (ML) and white-box simulation using the Synchronous Advantage Actor-Critic (A2C) Deep Reinforcement Learning (DRL) algorithm and PalaCell2D for simulating epithelial sheet growth [19, 20]. The approach aims to enhance the final cell count and geometrical configuration, assessing the feasibility and relevance of various computational DSE methods in biofabrication.

The paper further details the state of the art (section 2), methodology (section 3), validation and results (section 4), and future research directions (section 5).

## 2. Background

In the study of biological complexity, it’s vital to connect protocols with their biological mechanisms to improve performance and explain results biologically. White-box models, which are preferred for this, face challenges with increased complexity and simulation time. Various white-box modeling methods exist, encompassing continuous, discrete, and hybrid types [21, 22]. Ordinary Differential Equations (ODEs) are common in systems biology [23], while Agent-Based Models (ABMs) and discrete models, though less standardized, offer significant potential in computational biology [24, 25, 26]. Petri Nets (PN) models stand out for their accessibility, completeness, and suitability for complex biological processes, including ontogenesis and host-microbiota interactions [27, 28, 29, 30, 31]. Additionally, vertex models are instrumental in capturing dynamics in epithelial sheets [32].

OvS approaches blend simulation with optimization to explore model behaviors under varied parameter settings, aiding in the design of biofabrication processes [17]. They assist in generating simulated biofabrication protocols for lab testing, prioritizing informative experiments, and refining the process. There are two primary OvS categories [17]: (i) metamodel-based and (ii) model-based.

Metamodel-based OvS uses metamodels to estimate input-output relations, reducing computational time but at the expense of accuracy and explainability [33, 34, 35]. In biofabrication, this approach has optimized bioprinting and bio-ink/scaffold combinations, primarily focusing on non-living elements [36, 37, 38, 39, 40, 41]. Its use for living parts is limited, with some Deep Learning (DL) models predicting cellular responses [41, 42], but these lack desired explainability.

Model-based OvS relies on white-box models, offering higher explainability but at higher computational costs [17]. It addresses complex design spaces using heuristic and meta-heuristic optimization strategies [43]. However, its application in biomanufacturing is mostly confined to non-living components and structural aspects pre-maturation, using methods like Finite Elements Model (FEM) and Computational Fluid Dynamics (CFD) modeling [44, 45, 46, 47, 48, 49, 50, 51]. These models typically require metaheuristic algorithms for optimization due to their large state spaces [16, 15, 52]. A past study exemplifies a model-based OvS approach using GA for simulating and generating biofabrication protocols for epithelial monolayers [18]. The high computational complexity of these simulations necessitates a metaheuristic optimization approach. While effective, this method operates offline, limiting its adaptability to real-time biofabrication processes and discarding intermediate simulation data valuable for optimization. Alternatively, combining DRL with white-box simulations addresses the limitation of large state spaces. Reinforcement Learning (RL), a ML method, self-learns through environment interaction, guided by numerical rewards [53, 54]. DRL, integrating DL, is well-suited for high-dimensional challenges like biofabrication, offering continuous learning and adaptability for real-world manufacturing optimization [55, 16].

## 3. Materials and Methods

This work proposes a computational DSE methodology where a RL model and the A2C algorithm learn optimal biofabrication protocols to optimize simulated biofabrication. Fig 1 presents the implementation of the proposed methodology.

**Figure 1:**
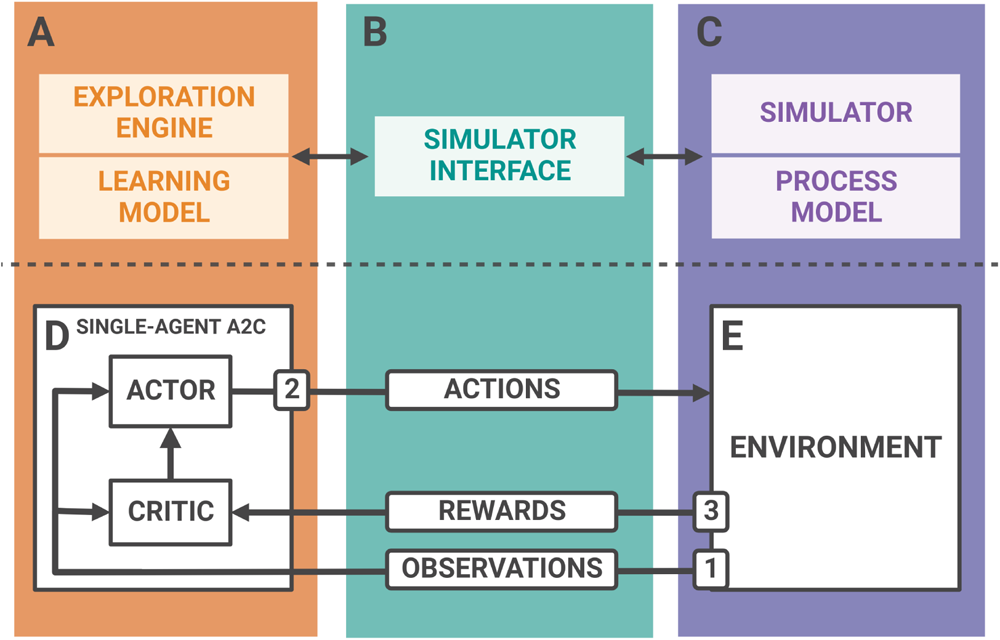
The components of the proposed framework and their interactions during the training process. (A) The exploration engine interrogates the learning model at each epoch. (B) The simulator interface allows the exploration engine to use the simulation performed by the simulator on the process model as a learning environment. (C) The simulator simulates the process model to simulate the application. (D) The proposed single-agent variation of the A2C algorithm interrogates the underlying ANNs, which generates distributions of actions to learn the best policy for a specific application based on the target of the optimization process through the interaction between the Actor and Critic. During each epoch, the optimization process flow includes these three steps, mediated by the simulator interface: (1) the communication of simulation states as observations from the learning environment to the learning model through the exploration engine; (2) the communication of a chosen action generated by the A2C algorithm by interrogating the underlying learning model; (3) the communication of rewards from the learning environment to the Critic in the learning model to evaluate the chosen action to further the training process.

The exploration engine (Fig 1.A) relies on a single-agent variation of the A2C algorithm and an Artificial Neural Networks (ANNs) as a learning model. The training process occurs through the interaction between Actor and Critic, and learning rules and rewards adapt to the particular application (Fig 1.D). The simulator interface (Fig 1.B) allows the exploration engine to interact with the simulator of the process model. The simulator of the process model (Fig 1.C) works as a learning environment (Fig 1.E) for the training process. The training process consists of epochs. Each learning epoch aims to generate the best possible action to take the learning environment closer to the target and contains the three following steps (numbered in Fig 1).

1. **Observation collection**: the exploration engine reads the simulation state as an observation from the learning environment, passing it to the learning model.
2. **Action generation**: the learning model, based on the received observation, generates an action that the exploration engine sends to the simulator for it to perform in the learning environment.
3. **Reward collection**: the exploration engine reads the reward associated with the performed action from the learning environment. It passes it to the Critic in the learning model to evaluate the effect of the chosen action in regards to reaching the goal of the optimization process Fig 1.C.

Each epoch results in a generated action. At the end of the training process, the sequence of generated actions to reach the optimal target defines an *optimal biofabrication protocol*.

The following subsections provide a detailed description of the exploration engine (subsection 3.1), the learning model (subsection 3.2), and the simulator interface (Appendix C), which are the constitutive components of the proposed methodology. Since they are application-specific, the section 4 provides a more detailed description of the simulator and the process model. Finally, subsection 3.4 illustrates the training process.

### 3.1. Exploration engine

This work relies on a RL model for learning. RL aims to find an optimal agent behavior to get maximal rewards. In particular, the exploration engine relies on Actor-Critic algorithms [56], which combine Q-learning [57] and Policy Gradients [58]. Policy gradient methods directly model and optimize a policy, representing an optimal agent behavior. In Actor-Critic, the Actor component generates an action to perform over the learning environment by random sampling from the probability distribution of all performable actions in a given state, relying on the same objective function as the *REINFORCE* algorithm [59].

The policy *π* is usually modeled with a parameterized function for the parameter vector *θ*: *π_θ_*(*a|s*). The value of the reward *J* (*θ*), which works as the objective function, depends on the chosen policy (see Equation 1). In Equation 1, *d^π^*(*s*) is the stationary distribution of Markov chain [60] for *π_θ_*, *a ∈ A* are actions, *s ∈ S* are states, *V ^π^*(*s*) and *Q^π^*(*s, a*) are, respectively, the value of state *s* and the value of a (*s, a*) pair, when following a policy *π* [61].

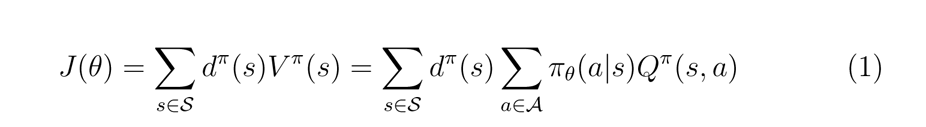

The goodness of the performed action *a* is evaluated based on the reward *J* (*θ*) generated by its execution in the learning environment. According to the *Policy Gradient Theorem* [62], the derivative of the expected reward is the expectation of the product of the reward and gradient of the log of the policy *π_θ_*. This allows us to compute the objective function without considering the derivative of the state distribution *d^π^*(*s*), untying calculations from the double dependency on action selection and states distribution [61]. In other words, the state distribution depends on the policy parameters, but there is no dependency of the policy gradient on the gradient of the state distribution [63]. This simplifies the calculation of the gradient *∇_θ_J* (*θ*) dramatically, as in Equation 2.

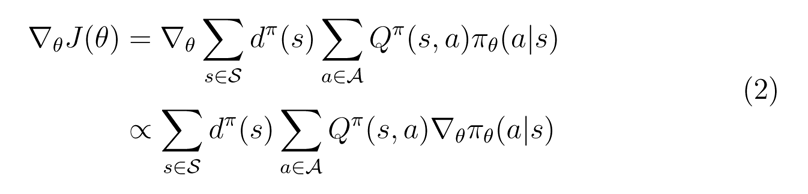

In Actor-Critic algorithms, the Critic component evaluates the goodness of the performed action *a* computing the reward *J* (*θ*) as the Mean Squared Error (MSE) between the current value and the best possible action [63], transforming Equation 1 into Equation 3, where *T* is the last timestep.

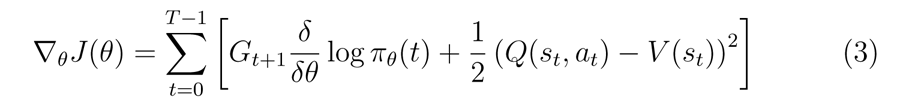

Since random sampling from action distributions can determine high variability in the log probabilities generated, letting the policy converge to a sub-optimal choice, the A2C algorithm [19] adjusts the log probabilities by multiplying them by the advantage. The advantage is a factor that quantifies how better the chosen action compares to other possible actions (Equation 4).

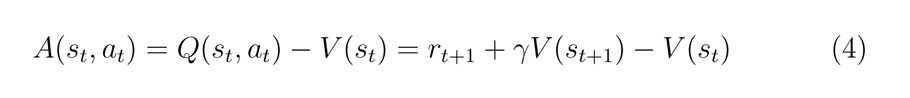

By applying Equation 4 within Equation 3, we obtain a more stable training process, as depicted in Equation 5, where *A*(*s_t_, a_t_*) is the advantage.

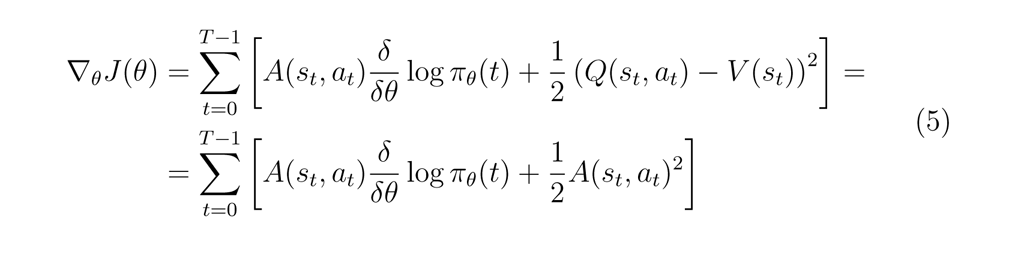

The exploration engine applies the actions generated by the ANN model in the environment. Then, after the effect of such actions, it generates the values to compute the A2C objective function, which in turn guides the update of the ANN weights through backpropagation [64].

Synchronous A2C [64] devises more agents acting simultaneously and combines the results they independently obtain, resolving inconsistencies. Each agent is a different ANN model, comprising an Actor and a Critic component as detailed in section 1.

To balance the vast capabilities of DRL with the computational feasibility of simulation-based DSE, this work proposes a simplified variant of the A2C algorithm where learning relies on a single agent. Overcoming the drawback of synchronizing multiple agents has a cost in accuracy. However, it allows to speed up the interaction with the environment while still expressing the capability to exploit response information and dynamically adapt to the system evolution.

### 3.2. Learning model

The learning model leverages the general ANN structure devised by the A2C algorithm [64]. The ANN structure includes a backbone (Fig 2.A) that processes the simulation states as observations and provides its outputs as an input to both the Critic and the Actor components. The ANN model implementation relies on the tensorflow library (version 2.9.1) [65], while the backbone relies on the ResNet18 model [66].

**Figure 2:**
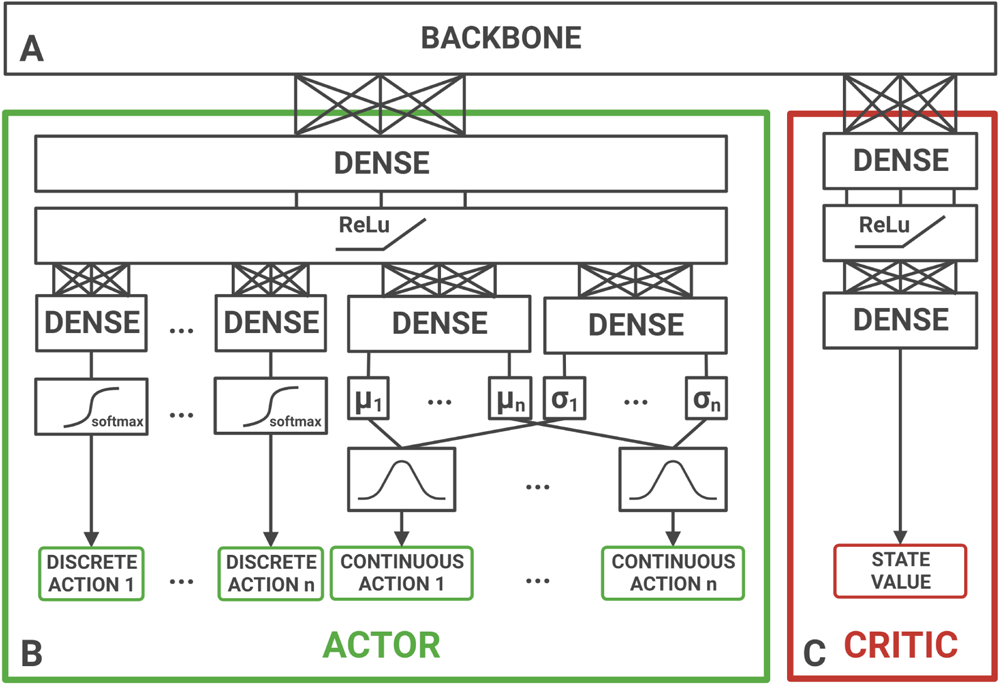
Actor-Critic ANN model. (A) A single backbone processes observations from the learning environment to feed the Critic and the Actor. (B) The Actor processes the backbone output to generate discrete and continuous actions over the learning environment. The architecture ANN adapts to the type and number of actions to generate. For discrete actions, the number of parallel output layers (based on softmax activations) equals the number of discrete actions to be generated (in the Figure, ranging from 1 to *n*). Each layer has an output size corresponding to the number of possible values for the discrete action it handles. For continuous actions, the number of action distributions in output and the number of neurons in both mean (*µ*) and variance (*σ*) layers equals the number of continuous actions to be generated (in Figure, ranging from 1 to *n*). Distributions support sampling of the continuous value for each of the actions considered. (C) The Critic processes the backbone output to generate the current state value.

#### 3.2.1. Model structure

The Actor (Fig 2.B) processes the backbone output to generate actions. It can produce both discrete and continuous actions. In particular, for each discrete one, the Actor generates a single value among the possible ones, leveraging a softmax function. The Actor component clips the output values of variance layers and discrete action layers between 10*^−^*^2^ and 10^3^ to avoid numerical issues in computation during the training phase. On the other hand, for each continuous one, it generates two values: (*µ*, the mean, and *σ*, the variance of a Gaussian distribution from which it randomly samples a value for generating the action. The Critic subnetwork (Fig 2.C) processes the backbone output to generate the current state value.

### 3.3. Model adaptability

The ANN model adapts to the application and training process. The Actor and Critic components have a fully connected Multi-Layer Perceptron (MLP) as an input layer. The number of input neurons in such layers is a hyperparameter for the training process. The size and shape of data provided to the ANN as an input determine the size and shape of its output layers. The width and height of the image-like observation of the environment, and channels, which is the number of color-like channels, set the input shape of the ANN.

The ANN output size adapts to the set of actions to generate, ranging from 1 to *n* in Fig 2, for both continuous and discrete actions. The number of continuous actions to generate sets the number of action distributions in output and the number of neurons in both the mean (*µ*) and the variance (*σ*) layers. Distributions support sampling of the continuous value for each of the actions considered. The set of possible values for discrete actions sets the number of parallel output layers with softmax activations.

### 3.4. Training process

The training process, described in Algorithm 1, relies on the train class, which interacts both with the environment and the model, implementing the A2C algorithm. The class manages training parameters, such as the learning rate (lr) and the discount rate (gamma), and supports getting relevant information in real-time and setting training length and starting epoch, allowing the restoration of the training from a previous state through the saved data if needed.

At first, the training creates the ANN model with the required input size and its optimizer (Algorithm 1, line 1). If restoring a previous state, it loads checkpoint files. Then, it instantiates the learning environment (Algorithm 1, line 2) and checks its integrity. The training loop continues until it reaches the maximum epoch number (Algorithm 1, line 3). The loop includes data collection through a GradientTape tensorflow section to track the data flow in the model. It is possible to show information in real-time during the training loop. Each epoch in the training loop consists of six main steps, operated by the train method (Algorithm 1, line 4-10).

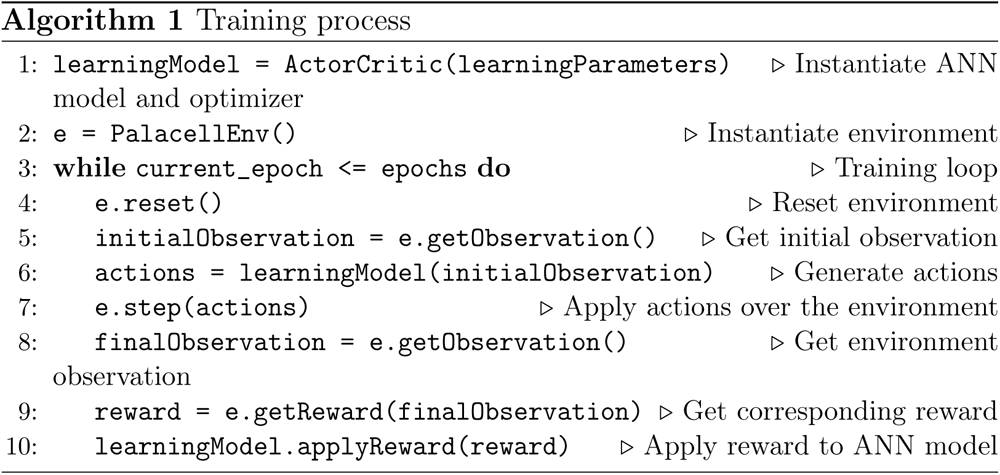

The first step (Algorithm 1, line 4) prepares the environment for the next learning episode, starting from the new initial state, by calling the reset method (Algorithm 1, line 4), and getting the first observation (Algorithm 1, line 5). The learning model uses the collected observation to generate actions (Algorithm 1, line 6). It also generates the probability of getting the obtained action and the state value that will support the loss computation at the end of the epoch. The step method (Algorithm 1, line 7) applies the generated actions on the learning environment. After that, the process gets a new observation of the learning environment, capturing its new state after the execution of the actions, and collects the corresponding reward (Algorithm 1, line 8-9). The training feeds the rewards into the learning model for learning (Algorithm 1, line 10). The Q-values and advantages, the saved state value, and the probability of getting the obtained action allow computing the objective functions defined by A2C for both Actor and Critic losses. The two losses are summed together and used by the GradientTape section as the loss to compute the gradients for backpropagation in the ANN through the optimizer.

The train_manager class allows implementing different environments for different sets of hyperparameters to perform distinct parallel training processes (Fig 3.A) *via* its parallel_train method, relying on the process class from the Python module multiprocessing, and a pipe, that is, a communication channel, passed to each subprocess. Each subprocess contains a whole and independent training process, based on the interactions between the train class (Fig 3.B), the environment file implementation (Fig 3.C), and the learning model (Fig 3.D), that allow the individual A2C agent (Fig 3.E) to learn by iteratively interacting with the learning environment (Fig 3.F). Finally, each training subprocess is stored so that it is possible to request real-time information through the pipe and wait for all the subprocesses to end their training.

**Figure 3:**
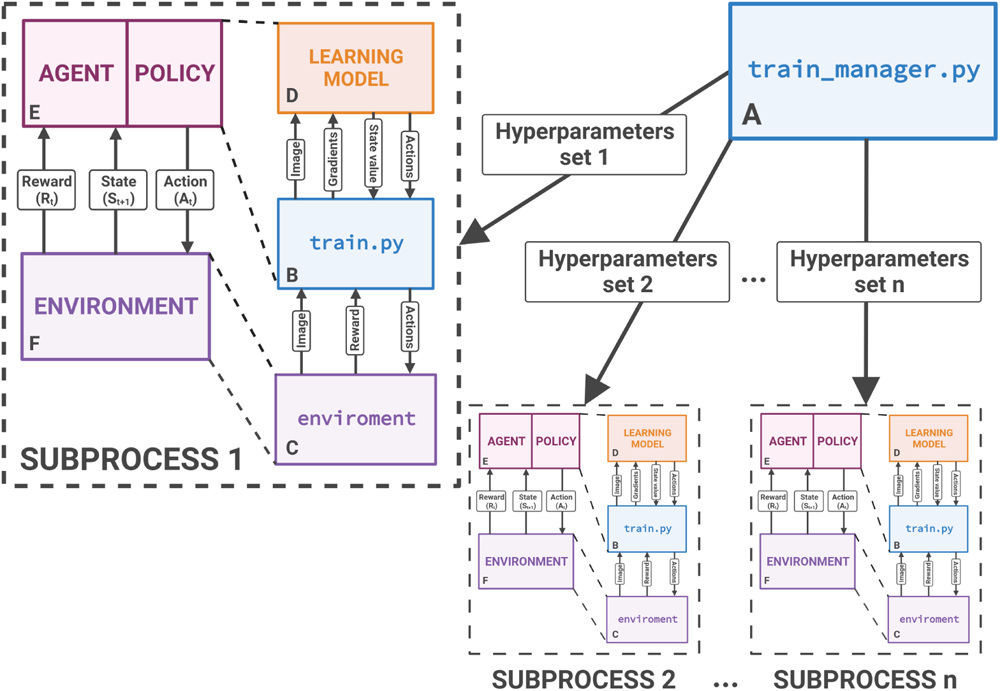
The proposed implementation of the A2C learning algorithm to support multiple training processes running in parallel. The train_manager (A), which consists of a single function, parallel_train, runs multiple training processes, each one with a different hyperparameter set, in parallel. Each subprocess contains a whole and independent training process based on the interactions between (B) the train class, (C) the environment file implementation, and the (D) learning model, that allows the individual A2C agent (E) to learn by iteratively interacting with the learning environment (F).

## 4. Results and Discussion

The proposed methodology is intended to perform DSE of protocol generation for real-world applications and physical biofabrication processes. In perspective, the optimization engine will interact with an automatic culture system, whose specific effectors define tunable parameters forming a biofabrication protocol. A first step towards real-world applications is optimizing protocols for simulated biofabrication.

To obtain meaningful results, this work relies on PalaCell2D, a simulator of the proliferation of epithelial cells oriented to tissue morphogenesis [20]. The PalaCell2D setup allows simulating tissue growth under different compression stimuli (see subsection 4.1). The optimization engine must communicate with the biofabrication process to control these stimuli, which will require interaction with a bioreactor for real-world applications. To this aim, an implementation of a simulator interface (see Appendix Appendix C) supports the interaction between the optimization engine and the PalaCell2D simulator. Consequently, a protocol consists of a sequence of values for the PalaCell2D tunable parameters. Eventually, the generation of optimized protocols (see subsection 4.2) aims to find the best sequence of parameter values to optimize the final product at the end of the simulation.

### 4.1. The PalaCell2D environment setup

PalaCell2D relies on a vertex model [32] that represents cells through their membrane, modeled as a set of vertices (Fig 4). PalaCell2D simulations iteratively evaluate the vertice positions for each cell and compute the forces exerted on each vertex.

**Figure 4:**
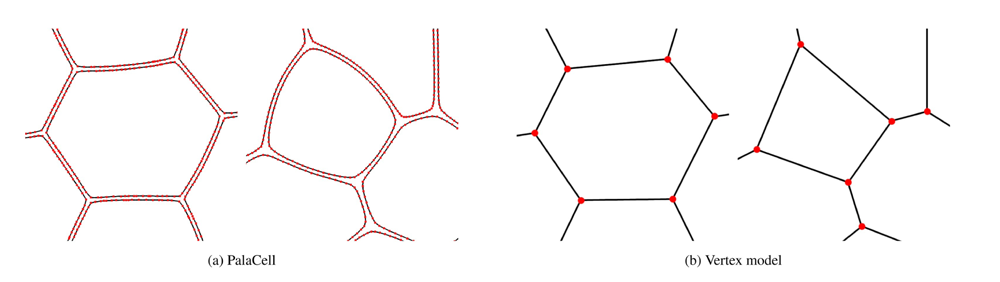
PalaCell2D models cell membranes as sets of vertices (the red dots) (a), basing on the vertex model approach (b) (taken from Conradin et al., 2021 [20]).

In the PalaCell2D model, the number of vertices that model a cell changes dynamically with its size so that the density of vertices remains uniform along the cell membrane. It supports the simulation of (i) mechanical properties, (ii) internal and external forces from the cellular, and (iii) extracellular compartments over the membrane. PalaCell2D aids in simulating cellular processes such as apoptosis and proliferation. Simulation controls them based on the internal cellular pressure. The latter depends on defined model parameters set for the simulation (as in Equation 6): the cell mass *m*, the cell area *A*, the pressure sensitivity *η*, and the target density *ρ*_0_.

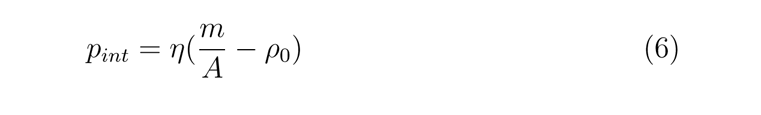

Internal pressure controls the growth of the cell mass. The mass changes at a rate that depends on the external pressure *p_ext_* applied on the cell, the target area *A*_0_, the mass growth rates during proliferation *ν* and relax *ν_relax_*, the simulation time step *dt* and the pressure threshold *p_max_* above which the proliferation stops as in Equation 7.

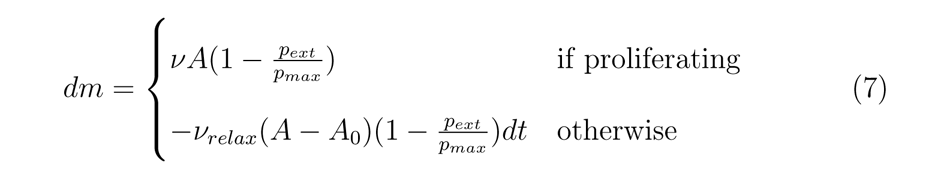

External pressure *p_ext_* depends on the contact with other cells and the external force *F_ext_*. The latter is either local (i.e., the interaction of a cell with a wall of the culture environment) or global (i.e., the force exerted over the whole culture environment).

The simulation scenario used for validation controls cell proliferation by applying a global external force (in Eq (8)). The parameter *a_prolif_*indicates the probability of controlling the switch to the proliferation state in cells.

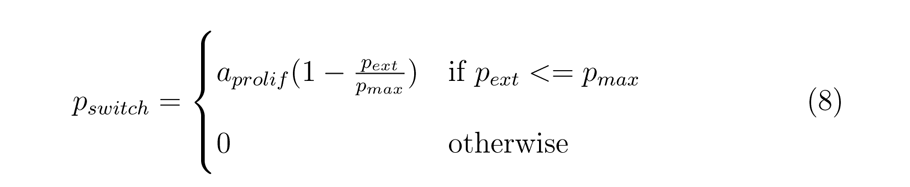

In the experiment proposed in [20], *F_ext_* models an external pressure exerted on a deformable capsule that acts on the cellular vertices with an intensity that depends on its distance from the center of the capsule and the radius of the capsule.

### 4.2. Optimized protocol generation

The capability of the DSE is assessed by demonstrating the framework capabilities to generate optimized protocols to obtain two different targets: (1) the maximal number of cells (see subsection 4.3) and (2) a precise shape of cells, i.e., a circular patch of cells (see subsection 4.4). Both targets are relevant for the biofabrication of epithelial sheets. The first includes maximizing the number of cells obtained, supporting the cell density required by a biomimetic TERM product to replace epithelia [67]. The second introduces the element of spatial control, aiming to optimize cell density and the epithelial sheet’s position and spatial organization within the culture system, still maximizing the number of cells. Moreover, the latter is a first step towards controlling the supracellular architecture, one of the significant open challenges in TERM biofabrication [68].

As stated before, the quality of the generated protocols depends on their ability to generate the target products at the end of a simulation. Protocols control parameter values representing the initial positioning of cells and the compression stimuli along the simulation (see Appendix B). Thus, they have two sections:

1. the initial section sets the value of the initialPos parameter once;
2. the second section sets the value for the comprForce and compressionAxis parameters at each simulation step.

Within this structure, the framework aims to learn the values for each parameter in an optimal protocol. Objective-specific metrics measure the similarity of the protocol-based final product to the target. Such a similarity is the keystone to protocol performance. The training process evaluates and saves metrics every 5 epochs to analyze training performance.

The following sections describe the experiments and their results, including learning performance along the training process and optimal protocols generated. These show the impact of different combinations of hyperparameters on learning performance. Since an exhaustive DSE of the target biofabrication processes is unfeasible in finite computational times [69], the results aim to analyze the evolution of learning processes to reach optimality rather than its convergence to a global optimum.

### 4.3. Target 1: maximization of the final number of cells

The first target aims to maximize the number of cells within the epithelial sheet at the end of a simulation. The tunable parameters in the protocol are compressionAxis and comprForce, and the reward value at the end of each epoch is the increment in the number of cells compared to the previous epoch. In the framework setup (see Appendix B), the simulated space in the environment is a 400x400x1 grid. The learning process targets compression stimuli and not the initial position of the cell, which holds constant (initialPos=(x=200, y=200)). This section analyzes the performance of the proposed approach by exploring different combinations of learning (see subsubsection 4.3.1) and simulation (see subsubsection 4.3.2) hyperparameters.

#### 4.3.1. Exploring lr and gamma learning hyperparameters on Target 1

This section analyses the performance of different combinations of learning hyperparameters: the *learning rate* (lr) and the *discount rate* (gamma).

Each experiment runs an independent training process based on a different combination of lr and gamma. Each training process includes 70 learning epochs. Each epoch corresponds to a simulation with a total number of 3400 simulation steps. During each epoch, the optimization engine interacts with the simulator every 20 (numIter=20) simulation steps, while performance evaluation and saving occurs every 5 epochs. Tab 1 illustrates the six combinations explored, summarizing the highest absolute final number of cells obtained as a performance measure for every setting.

**Table 1:** Highest final number of cells for Target 1 for the lr **and** gamma **learning hyperparameters exploration.** For each experiment, the last column reports the highest final number of cells obtained across training.

| Experiments | Hyperparameters |  | Highest final number of cells |
| --- | --- | --- | --- |
|  | lr | gamma |  |
| *Exp1* | 0.001 | 0.99 | 270 |
| *Exp2* | 0.001 | 0.95 | 269 |
| *Exp3* | 0.0001 | 0.99 | 267 |
| *Exp4* | 0.0001 | 0.95 | 266 |
| *Exp5* | 0.00001 | 0.99 | 267 |
| *Exp6* | 0.00001 | 0.95 | 268 |

Fig 5 visualizes learning performance as the distribution of the final number of cells obtained within six sliding windows of 20 learning epochs (0-20, 10-30, 20-40, 30-50, 40-60, and 50-70, respectively). Violin plots in Fig 5 depict the distributions of these values with probability density curves whose width represents the frequency of data points in each value range inferred from the data. These are augmented by boxplots, which detail the lower and upper quartiles at the box’s ends and the median value inside the box. Additionally, the lines stretching vertically from the box reach up to the maximum non-outlier values, which are the data points lying within a distance no more than 1.5 times the interquartile range from the box’s edges [70].

**Figure 5:**
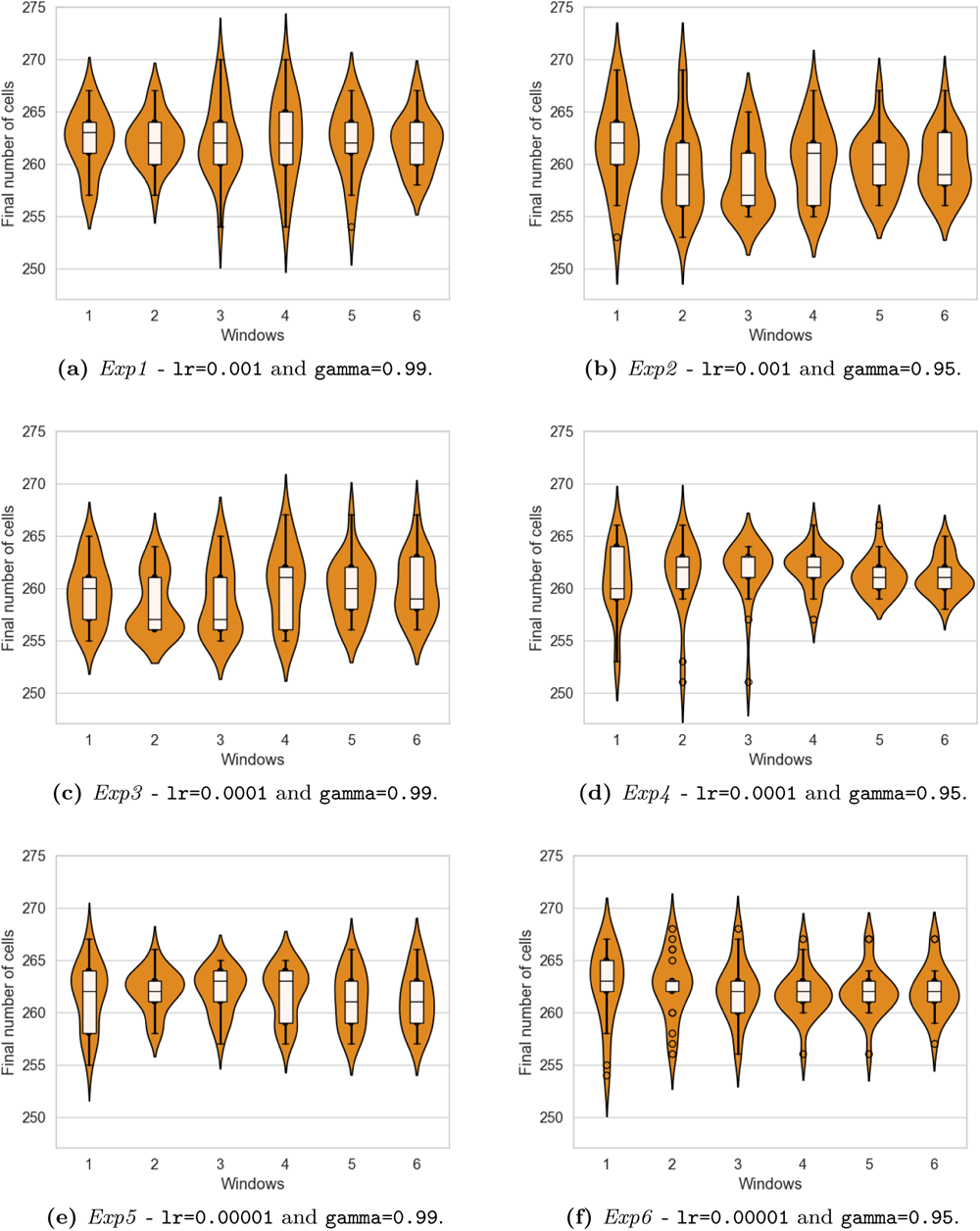
RL learning performances on Target 1 - learning hyperparameter values exploration. Each panel shows the performance of one training process over six windows of 20 learning epochs, based on a different combination of the lr and gamma learning hyperparameters.

Results show that the DSE alternates exploration and exploitation along learning phases, exhibiting a range of tendencies with different combinations of lr and gamma.

- *Exp1* (lr=0.001, gamma=0.99, Fig 5a) is the experiment yielding the highest number of cells (270, see Tab 1). This process starts by exploring the upper portion of the value range at window 1. Its learning performance marks a peak at window 4, where the upper quartile extends to a higher value range, and the upper whisker marks the peak value. Subsequently, performance lowers, and the learning explores smaller value ranges.
- *Exp2* (lr=0.001, gamma=0.95, Fig 5b), while yielding a similar highest value (269, see Tab 1), exhibits an overall broader distribution of fitness at window 1, compared to *Exp1*. There is a remarkably higher variability among median values across windows, and several violin plots exhibit probability density peaks in the lower range of fitness values (windows 2, 3, and 6, respectively).
- *Exp3* (lr=0.0001, gamma=0.99, Fig 5c) shows remarkable inter-window variability of both median values and density profiles, with window 2 showing an evident bimodality with two peaks at 256 and 261, respectively. This marks an exploration phase for the learning process, which reaches higher performances starting at window 4, corresponding to the broadest density profile and the highest median value along the training. Finally, across the last windows, this training exhibits a stable increase in the mean performance and a stable decrease in its variance, marking the training exploiting knowledge to find a local optimum.
- *Exp4* (lr=0.0001, gamma=0.95, Fig 5d) shows pronounced exploration starting from the first window, where the distribution of values spans the middle and lower ranges. In the following two windows, the yield of outlier values extends to the bottom part of the range, as indicated by the long tails of the density profiles combined with the reduced size of the boxplot bottom whisker. Then, the learning focuses on a limited range of values, which still yields modest and decreasing performances.
- *Exp5* (lr=0.00001, gamma=0.99, Fig 5e) starts by exploring a broad value range at window 1 to start then to exploit knowledge at window 2, showing a compact interquartile range and a density profile peak around 263. Then it gradually progresses towards increasing medians and, at the same time, broader explored ranges.
- *Exp6* (lr=0.00001, gamma=0.95, Fig 5f) has a peculiar behavior: while most windows correspond to compact interquartile ranges, they also mark the presence of several outliers. This process generally marks a passage from an initial exploration phase to a steady, lower performance level.

Fig 6 shows the impact of the optimization process, visualizing the structure of the protocols generated at the beginning of training at epoch 0 (Fig 6a) and at epoch 37 (Fig 6b)) of *Exp1* (lr=0.001, gamma=0.99). When no optimization has occurred, the protocol obtained at epoch 0 exhibits a disorganized structure. The protocol obtained at epoch 37, after training advanced, which yields the maximum number of final cells across experiments (270), keeps the stimulation in the very low range between comprForce=0.005 and comprForce=0.0055 over a single compression axis (compressionAxis=’X’). Each point in the plot marks a learning episode in which the optimization engine interacts with the simulator, setting new values for stimulation parameters comprForce and compressionAxis. Each learning episode controls numIter=20 steps in the simulation. The x-axis represents the sequence of 170 learning episodes along a simulation of 3400 simulation steps. A low-compression protocol is consistent with the target of maximizing the final cell number, according to the knowledge of the process and PalaCell2D modeling, that assumes the inverse relationship between external force at a cell and the probability the cell proliferates [20].

**Figure 6:**
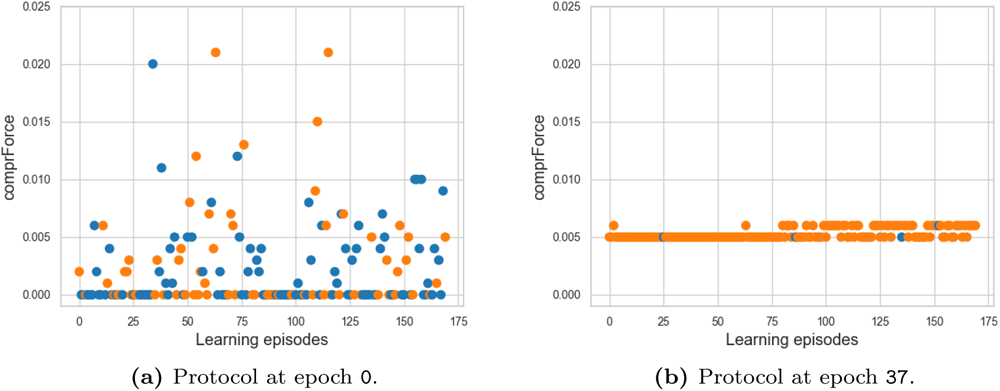
Compression stimuli protocols generated for Target 1 at epoch 0 and epoch 37 during *Exp1* (lr=0.001, gamma=0.99). Dots represent the stimulation at each learning episode, including the set comprForce compressionAxis values, either ’X’ (orange dots) or ’Y’ (blue dots). Each learning episode corresponds to one interaction of the learning process with the simulator, setting new values for stimulation parameters comprForce and compressionAxis for numIter=20 simulation steps. The 170 learning episodes shown occur along a simulation of 3400 simulation steps.

#### 4.3.2. Exploring the numIter simulation hyperparameter on Target 1

Starting from the exploration of learning hyperparameters, this section further analyses the *Exp3* setup (lr=0.0001, gamma=0.99) by exploring different values of the numIter simulation hyperparameter, since this hyperparameters combination exhibits a good balance between exploration and exploitation (see subsubsection 4.3.1).

Tab 2 illustrates experimental results exploring three values of numIter: 20 (kept as a reference from the previous exploration), 50 and 100, alongside the highest final number of cells obtained for every setting.

**Table 2:** Highest final number of cells for Target 1 for the numIter **simulation hyperparameter exploration.** Each experiment corresponds to a fixed combination of the lr and gamma parameters and a different numIter value. The last column reports the highest absolute number of cells obtained during each experiment.

| Experiments | Hyperparameters |  |  | Highest N of cells |
| --- | --- | --- | --- | --- |
|  | lr | gamma | numIter |  |
| *Exp3* | 0.0001 | 0.99 | 20 | 267 |
| *Exp7* | 0.0001 | 0.99 | 50 | 274 |
| *Exp8* | 0.0001 | 0.99 | 100 | 278 |

Fig 7 visualizes learning performance as the evolving distribution of the values of the final number of cells obtained along training processes. Performance is evaluated within six windows of 20 epochs. Results show that increasing values for numIter improves the overall learning performance across experiments. Indeed, *Exp7* (numIter=50, Fig 7b) exhibits consistently higher mean values than *Exp3* (numIter=20, Fig 7a) along training. Median values lie above 265 across all windows in *Exp7*, while they stay below 261 in *Exp3*. Also, density profiles show peaks above 265 and very small interquartile ranges. Window 5 marks the sharpest range. *Exp8* (numIter=100, Fig 7c) shows even higher median values, starting higher than 270 and reaching 277 in the last windows. This training shows a clear exploitation with convergence to a local optimum, increasing performance and decreasing interquartile ranges after the second window. Results indicate a remarkable impact of the numIter simulation hyperparameter on performances, underlining that learning-simulation interaction frequency deeply affects learning dynamics.

**Figure 7:**
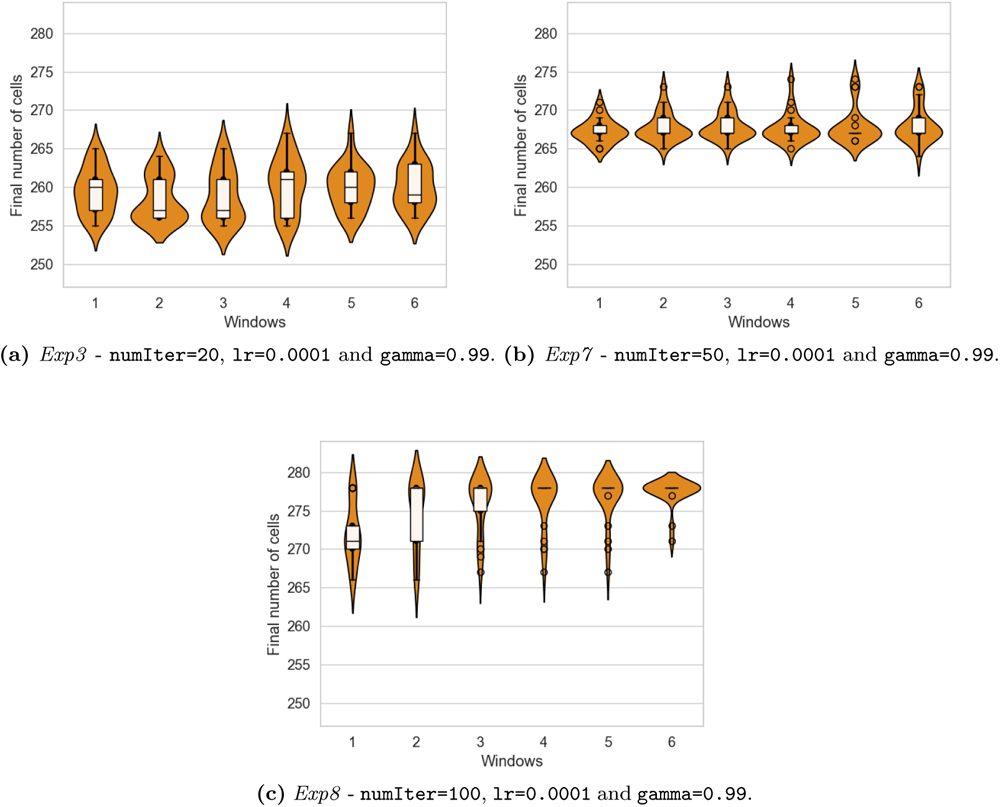
RL learning performances on Target 1 - simulation numIter **hyperpa-rameter values exploration.** Each panel shows the performance of one training process over six windows of 20 learning epochs, based on lr=0.0001, gamma=0.99, and different numIter values.

#### 4.3.3. Testing the ANN learning for Target 1

Compared with existing optimization approaches from the literature, one major strength of the proposed approach is its learning capability. This section provides the results of testing the models resulting from the three training processes presented in subsubsection 4.3.2: numIter=20 (*Exp3*), numIter=50 (*Exp7*) and numIter=100 (*Exp8*). Each testing process included 30 epochs. During testing, the framework maintains the functioning described for training, excluding the ANN learning step. The trained ANN model resulting from the training phase is loaded at the beginning of testing. When the framework interacts with the environment, it just leverages the trained ANN to generate actions and administer them to the simulator. It does not use the resulting observations to train the ANN further.

Fig 8 visualizes the distributions of the 30 different final numbers of cells obtained during testing epochs, exhibiting relations to learning performances. In particular, the ANN trained on *Exp3* (numIter=20) generates a broad distribution of fitness values ranging from 255 to 270, and the most frequent values range between 256 and 258, with a peak at 256 (Fig 8a). During training, *Exp3* explores this range of values across windows, as delimited by boxplot whiskers in Fig 7a). In addition, the central quartiles across windows lie in a similar range to the testing fitness value distribution. The ANN trained on *Exp7* (numIter=50) generates a way narrower distribution of values peaking at 267 (Fig 8b). This value is coherent to the narrow density profile peaks across windows in Fig 7b). In addition, the distribution tails encompass the range where the central quartiles lie across windows. Finally, the ANN trained on *Exp8* (numIter=100) generates a similarly narrow distribution of values peaking at 278 (Fig 8c)). Again, this value is coherent to the density profile peaks across the last three windows in Fig 7c), and the distribution spans a range that also characterizes values generated by training.

**Figure 8:**
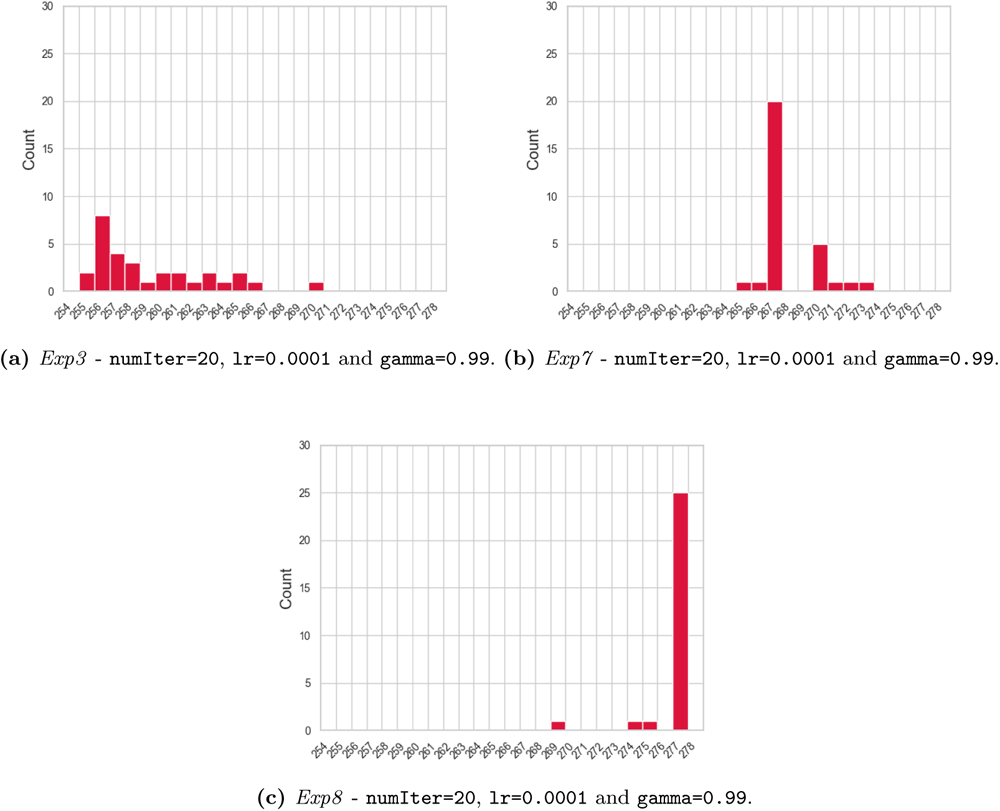
RL testing performances on Target 1 based on the numIter **hyperparameter values exploration.**. Histograms report distributions of the final number of cells obtained after each testing epoch. Each panel shows the performance of a trained model based on lr=0.0001, gamma=0.99, and different numIter values.

#### 4.3.4. Discussion on Target 1

The results collectively illustrate the effectiveness of the RL approach in optimizing cell proliferation within the PalaCell2D simulation environment.

In summary, results on Target 1 show that the proposed RL approach is capable of:

- maximizing the final number of cells at the end of a simulation;
- optimizing protocols of compression stimuli coherently with the target and the modeling assumptions;
- expressing a range of training behaviors and performances with different combinations of learning and simulation hyperparameters (lr, gamma and numIter);
- learning and retaining knowledge within the trained ANN models, whose testing recapitulates training performance.

In subsubsection 4.3.1, the combination of lr and gamma values influences how quickly the model learns and how it values immediate versus future outcomes. Indeed, the provided grid search of value combinations leads to different learning behaviors, with some combinations leading to faster convergence at the risk of instability or gradual, stable learning at the cost of slower convergence. This interplay between lr and gamma is critical for tuning the ANN’s performance to optimize outcomes in the simulated epithelial cell environment. Indeed, the choice of the *Exp3* setup (lr=0.0001, gamma=0.99) for the subsequent numIter exploration is based on the good balance between exploration and exploitation. The numIter parameter controls the interaction frequency between the learning algorithm and the simulation environment. In subsubsection 4.3.2, the lowest numIter=20 value, corresponding to more frequent updates of the learning strategy based on feedback from the simulation, leads to quicker but less stable learning dynamics (Fig 7a). On the other hand, higher numIter=50 and numIter=100 allow for more extended periods of simulation before each learning update, leading to a deeper understanding of the long-term consequences of actions at the cost of slowing down the learning process (Fig 7b and Fig 7c). The density profiles and boxplots in the second part of the training suggest that the model is overfitting when numIter=100. Limiting the number of interactions between the learning and the environment by setting the numIter hyperparameter can make the ANN excessively tailored to the training data, shifting from beneficial exploration of diverse data patterns to excessive exploitation, leading to high accuracy on training data but poor generalization to new, unseen data. Results clearly show that this balance between rapid adaptation and in-depth exploration directly impacts the learning stability and convergence quality, exploring the trade-off between immediate and future rewards in complex simulated environments. The ability of the ANN models to generalize their learning is evident from the testing results. When tested, each trained model produces outcomes consistent with its training process (see subsubsection 4.3.3), indicating that the models have effectively learned and internalized the patterns and dynamics of the simulation environment. Target 1 is intrinsically limited to the functional readouts from the environment, having the final number of cells as a measure of fitness while sustaining a first demonstration of framework capabilities.

### 4.4. Target 2: precise maximization of the fraction of cells within a circular area

Optimization in real-world applications often includes targeting culture processes’ functional and structural complexity. In this direction, Target 2 aims to maximize the cell fraction within a circular area, intending to fabricate a circular cell patch with a defined position and dimension at the end of a simulation. In this case, the maximization problem meets the goal of positioning the initial cell so that, proliferating, cells overlap a designated circular target space. The reward is the fraction of cells within the target (*fraction_inside_*), as in Eq (9) where *cells_inside_* and *cells_outside_* are the number of cells inside and outside the target space, and *cells_final_* is the total number of obtained cells.

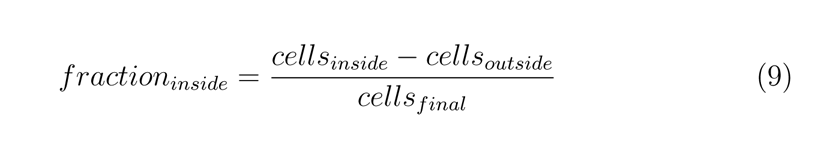

The *fraction_inside_* values along training measure performance. Protocols for this target control the initial cell position before the simulation starts and the compression stimuli along the simulation. The framework setup (see Appendix B) includes a simulated space based on a 400x400x1 grid, with the addition of a circular target area centered in x=200 and y=250, having a radius length of 80.

The learning process targets compression stimuli and the initial position (initialPos) of the first cell in the simulation. For all experiments, the exploration range for initialPos lies between 90 and 310 for both starting position coordinates. Thus, the generated protocols cover two steps:

1. initialPos parameter value setting (once, before the simulation starts);
2. comprForce and compressionAxis parameters values setting (along simulation).

This requires two learning processes to work together: the primary, outer training process learns initial positions, while a secondary, inner training process learns compression stimuli. Two learning environments, leveraging a hierarchy of subprocesses (see Appendix D) implement this design. The inner environment recapitulates the setup from subsection 4.3. Consequently, at each epoch, the outer training process generates a single action to set initialPos. Then, it launches the inner training process and waits for completion. During each outer epoch, the inner process undergoes training to learn compressionAxis and comprForce, interacting with the simulation every numIter=20 steps. Each inner epoch comprises a simulation of 2500 simulation steps. Each nested training process runs for 70 epochs, and performance evaluation occurs every 5 epochs.

#### 4.4.1. Exploring lr and gamma learning hyperparameters on Target 2

Each experiment runs an independent training process based on a different combination of two learning hyperparameters that affect both the outer and inner training: the *learning rate* (lr) and the *discount rate* (gamma). Tab 3 illustrates the six combinations explored, summarizing the highest absolute final fraction of cells obtained during training.

**Table 3:** Highest *fraction_inside_* **for Target 2 for the** lr **and** gamma **learning hyperparameters exploration.** For each experiment, the last column reports the highest *fraction_inside_* obtained across training.

| Experiments | Hyperparameters | | Highest $fraction_{inside}$ |
| --- | --- | --- | --- |
|  | lr | gamma |  |
| *Exp9* | 0.001 | 0.99 | 0.647 |
| *Exp10* | 0.001 | 0.95 | 0.642 |
| *Exp11* | 0.0001 | 0.99 | 0.475 |
| *Exp12* | 0.0001 | 0.95 | 0.638 |
| *Exp13* | 0.00001 | 0.99 | 0.000 |
| *Exp14* | 0.00001 | 0.95 | 0.640 |

Fig 9 illustrates the learning performance by showing the final fraction of cells across six sliding windows covering 20 epochs each (0-20, 10-30, 20-40, 30-50, 40-60, 50-70).

**Figure 9:**
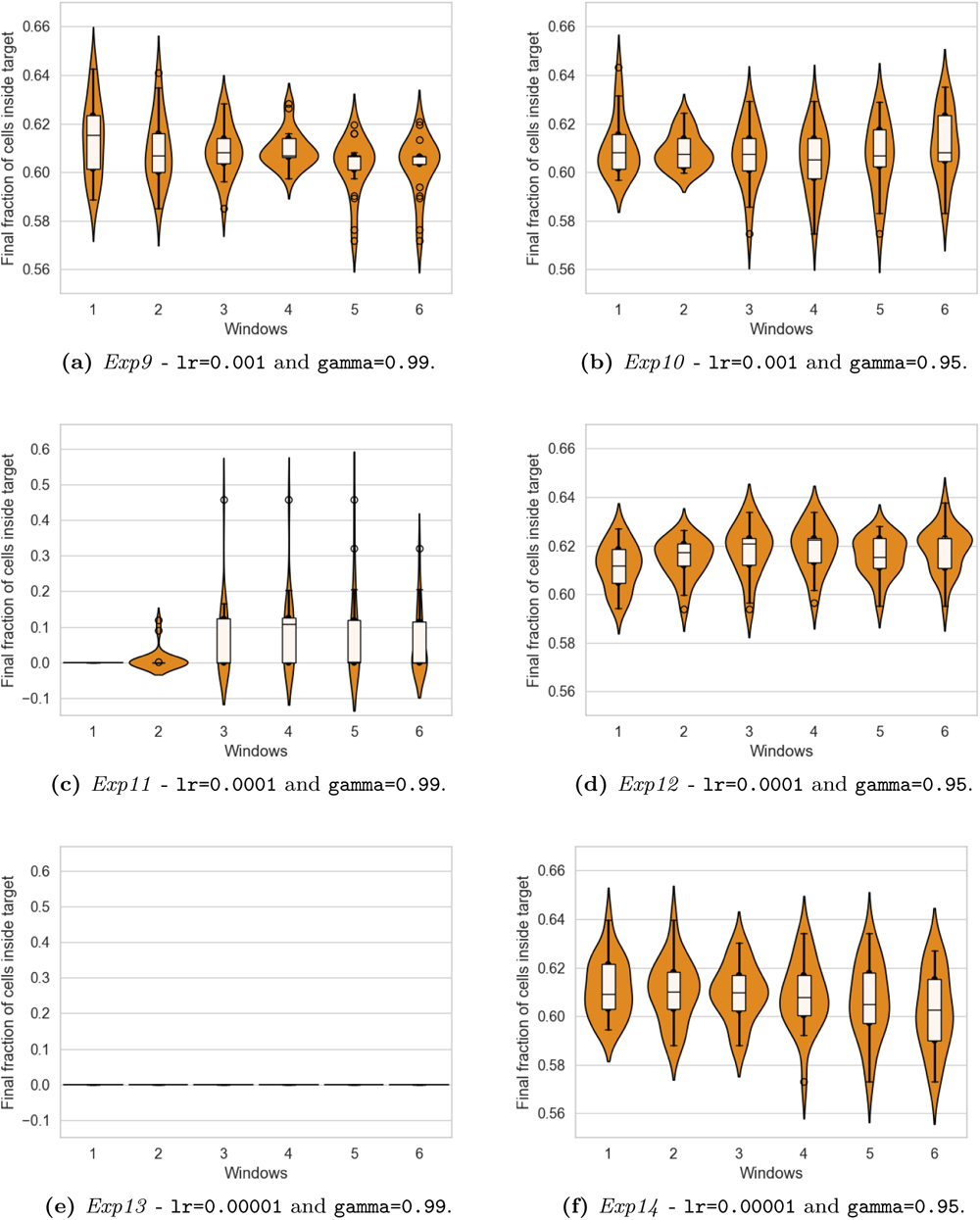
RL learning performances on Target 2 - learning hyperparameter values exploration. Each panel shows the performance of one training process over six windows of 20 learning epochs, based on a different combination of the lr and gamma learning hyperparameters.

Results show that the DSE alternates phases of exploration and exploitation along learning, exhibiting a range of tendencies with different combinations of lr and gamma.

- *Exp9* (lr=0.001, gamma=0.99, Fig 9a) yields the highest final fraction of cells across experiments (0.647, see Tab 3) within the first window, whose broad probability density profile and high interquartile range indicate a marked tendency towards exploration at the beginning. Profiles become increasingly compact in the fourth window, where learning stabilizes on a lower performance. This phase evolves in additional exploration during the last two windows, which exhibit reduced interquartile ranges, several fitness values marked as outliers, and broad density profiles, indicating marked exploration that is not converging to an optimum.
- *Exp10* (lr=0.001, gamma=0.95, Fig 9b) has a more stable dynamics, where after the initial two windows that generate values in a compact range centered over 0.61, the training enters an exploration phase characterized by broader profiles and increasingly wider interquartile ranges. While the median values stay centered around 0.61, the upper quartiles progressively extend towards higher values, approaching 0.62 at the last window.
- *Exp11* (lr=0.0001, gamma=0.99, Fig 9c) exhibits dramatically lower performances, with density profiles peaking between 0 and 0.1. Central windows include outliers spanning to the 0.4-0.5 range, which is still way lower than the ranges populated by the previous experiments (considering this, to maximize readability, the plot has a different scale for the fitness values).
- *Exp12* (lr=0.0001, gamma=0.95, Fig 9d) marks a tendency towards exploitation, where the training starts with a compact density profile peaking close to 0.62. Profile peaks gradually move towards higher values until reaching 0.62, while they consistently grow in width, indicating training is converging towards a local optimum.
- *Exp13* (lr=0.00001, gamma=0.99, Fig 9e)similarly to *Exp11*, has way lower performances. Still, in this case, the learning consistently yields null performance (again, the plot scale differs from the rest).
- *Exp14* (lr=0.00001, gamma=0.95, Fig 9f) illustrates the passage from an initial prevalence of exploitation with density profiles peaking around 0.61 towards increasing exploration, with profiles broadening to the lower ends of the value range, while the median stays close to the initial value across all training.

Fig 10 illustrates a bi-dimensional histogram of the X and Y coordinates of initialPos values generated by the training process based on the lr=0.0001 and gamma=0.95 setting (*Exp12*). Each panel corresponds to a learning window, covering the six windows from Fig 9d. The color intensity of squared sub-areas in purple grows with the total number of times the training chose a position within them during the window. The red circle indicates the target area, and its color intensity is directly proportional to the mean *fraction_inside_* in the window, normalized over the fitness value range for this experiment. The exploration range for initialPos lies between 90 and 310 for both coordinates, and this bounds the area where initial positions lie, with an offset linked to the bin size chosen to maximize figure readability.

**Figure 10:**
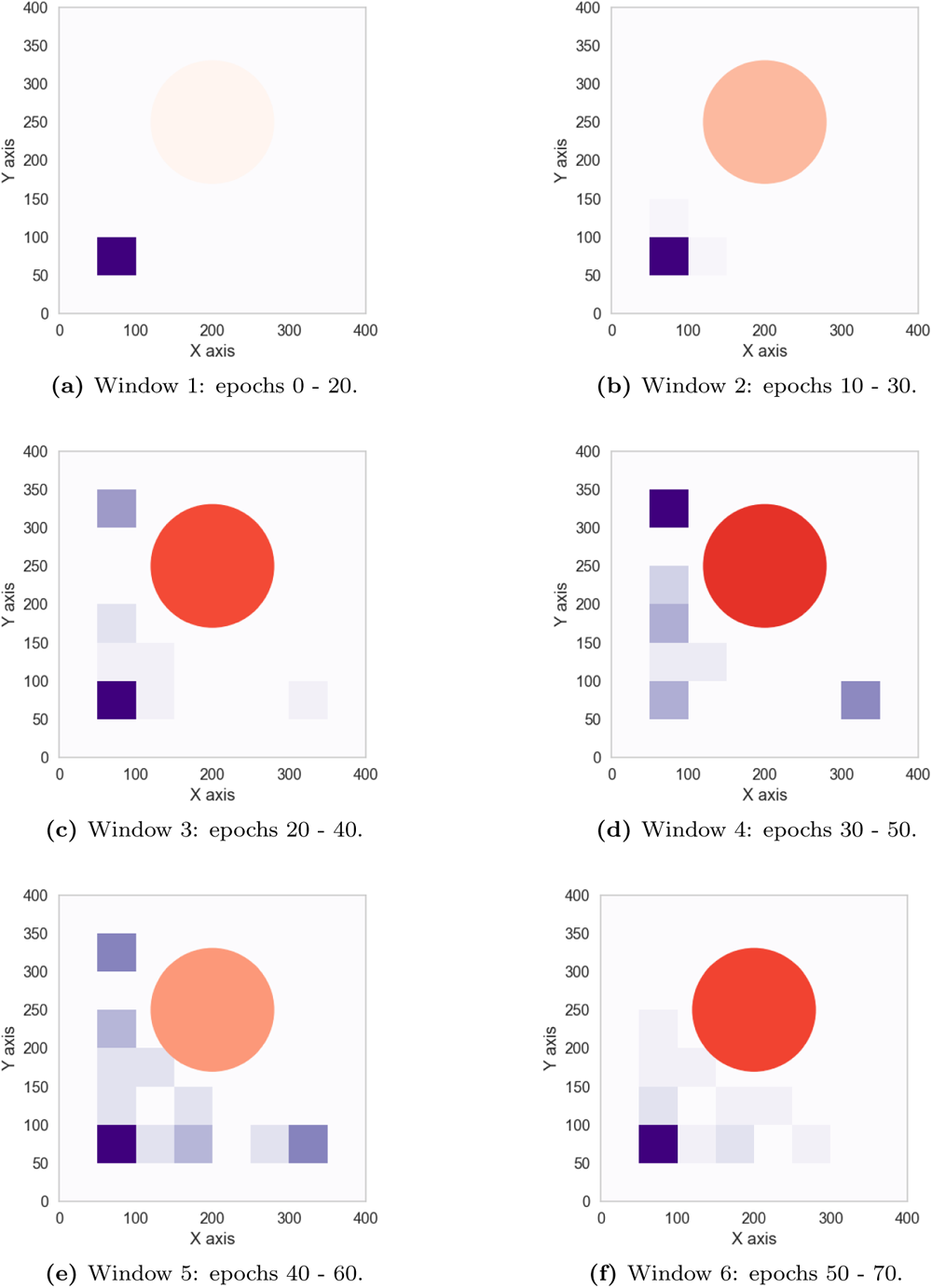
initialPos **coordinates generated for Target 2 based on** lr=0.0001 **and** gamma=0.95 **(*Exp12*).** The color intensity of squared areas represents the frequency as the initial position in 20-epochs windows within an exploration range between 90 and 310 for both coordinates. The red circle represents the target area, and its color intensity visualizes the normalized mean *fraction_inside_*.

During the first window (Fig 10a), the training exclusively generates initialPos value in the left bottom corner of the action space, which is distant from the target, resulting in the lowest normalized mean *fraction_inside_* across windows. The second window (Fig 10b) shows training starts to explore the adjacent positions to the bottom left corner, determining a slight increase in fitness. In the following windows, the training explores several initialPos coordinates closer to the target, resulting in higher normalized mean performance values, peaking at window 4 (Fig 10d). The latter shows that the most initialPos coordinates explored lie in the top left corner, which is close to the target, and thus sustains higher mean *fraction_inside_* for the corresponding epochs. On the contrary, windows 3 (Fig 10c), 5 (Fig 10e) and 6 (Fig 10f), while showing the same tendency to explore areas closer to the target, still have the majority of initialPos coordinates at the bottom left corner, which limits the performance of the corresponding epochs and lowers the mean *fraction_inside_*.

#### 4.4.2. Exploring the numIter simulation hyperparameter on Target 2

This section explores the impact of the numIter hyperparameter while learning hyperparameters hold constant values for lr (0.0001) and gamma (0.95) (from *Exp12*). This training yields a maximum *fraction_inside_* value of 0.638. Yet, this choice is not based on the highest performance value obtained across all training instances 0.647, which belongs to *Exp9* instead (see Tab 3). The choice is rather based on the fact this combination shows a steady and stable performance increase (see subsubsection 4.4.1).

Tab 4 illustrates the highest absolute fraction of cells obtained during each experiment. Results show performance is inversely proportional to numIter hyperparameter values. numIter=20, as illustrated in subsubsection 4.4.1, has 0.638 as the highest value. numIter=50 and numIter=100 yield highest values of 0.635 and 0.620, respectively.

**Table 4:** Highest final fraction of cells for Target 2 for the numIter **hyperparameter values exploration.** The last column reports the highest final *fraction_inside_* obtained across training for each experiment.

| Experiments | Hyperparameters | | | Highest $fraction_{inside}$ |
| --- | --- | --- | --- | --- |
|  | lr | gamma | numIter |  |
| *Exp12* | 0.0001 | 0.95 | 20 | 0.638 |
| *Exp15* | 0.0001 | 0.95 | 50 | 0.635 |
| *Exp16* | 0.0001 | 0.95 | 100 | 0.617 |

Fig 11 visualizes training performances over six windows, based on lr=0.0001 and gamma=0.95 learning hyperparameters values and different numIter values (from the top to the bottom): numIter=20 (*Exp12*), numIter=50 (*Exp15*), numIter=100 (*Exp16*).

**Figure 11:**
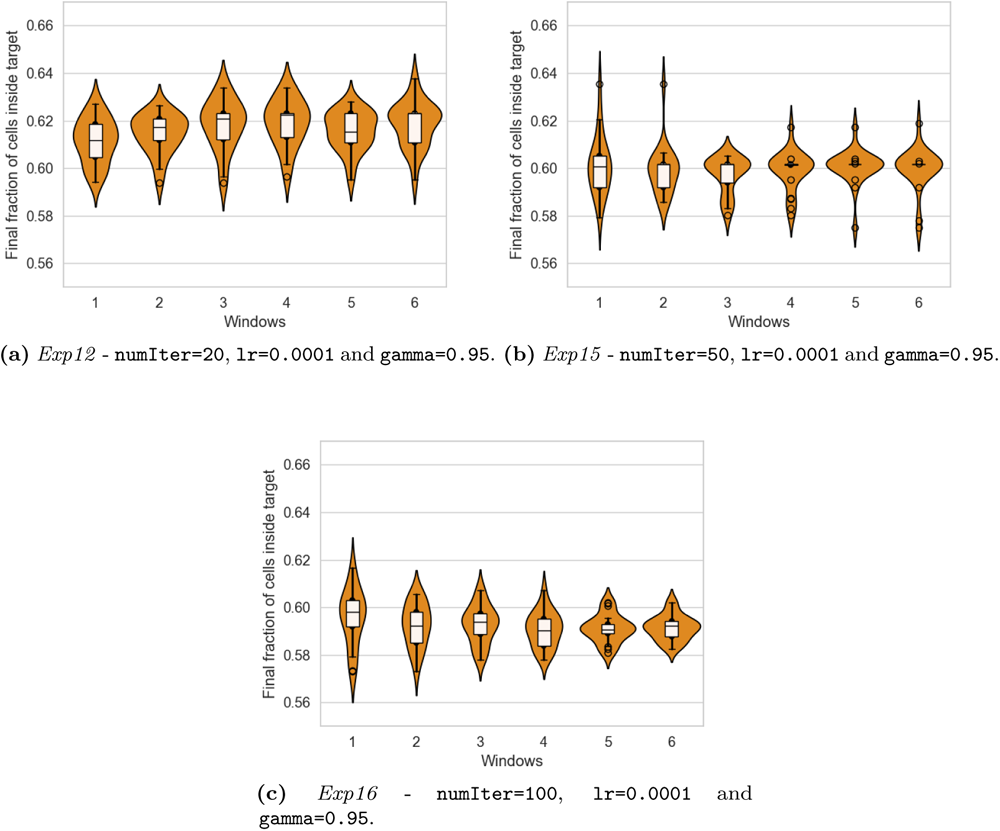
RL learning performances on Target 2 - simulation numIter **hyperparameter values exploration.** Each panel shows the performance of one training process over six windows of 20 learning epochs, based on lr=0.0001, gamma=0.95, and different numIter values.

Results show that numIter hyperparameter values visibly affect learning dynamics. Compared to *Exp12* (Fig 11a), based on numIter=20, *Exp15* (Fig 11b), based on numIter=50, shows probability density profiles with broader curves peaking at 0.60 (also the median value) in the first three windows. The second part of the training exhibits flattened interquartile ranges with a stable median at the same value, while several outliers broaden the density profile. This indicates the training is stabilizing over the initial performance, meaning the exploration did not sustain any improvements starting from there. *Exp16* (Fig 11c), based on numIter=100, shows that the training starts at a similar median value close to 0.60, with a broad density profile underlinging exploration. In the following windows, both density profiles and interquartile ranges shrink, centering their peaks and median values around 0.59, showing that exploitation yields lower performance than the one reached initially.

#### 4.4.3. Testing the ANN learning for Target 2

This section provides the results of testing the models from the three training processes presented in subsubsection 4.3.2. To obtain testing results, the framework leverages the trained ANNs to generate actions and administers them to the simulator, but it does not use the resulting observations to train them.

Fig 12 visualizes the distributions of the 30 values obtained for the final number of cells during testing epochs. As observed in subsubsection 4.3.3, testing results relate to learning performances. In particular, the ANN trained on *Exp12* (numIter=20) peaks at the interval between 0.61 and 0.62 (Fig 12a), and the distribution profile recapitulates density profiles across windows during training (Fig 11a). In addition, the distribution central domain encompasses the range where the central quartiles lie, and the right tail spans close to 0.64, as the upper whisker of the boxplot at window 6 does. The ANN trained on *Exp15* (numIter=50) generates a distribution of values whose two peaks lie between 0.595 and 0.605 (Fig 12b). This value is coherent with the narrow density profile peaks across windows in Fig 11b and with the median value training stabilizes onto. Finally, the ANN trained on *Exp16* (numIter=100) generates a distribution shifted towards lower values (Fig 12c) which are coherent to the density profile peaks across the last two windows in Fig 11c.

**Figure 12:**
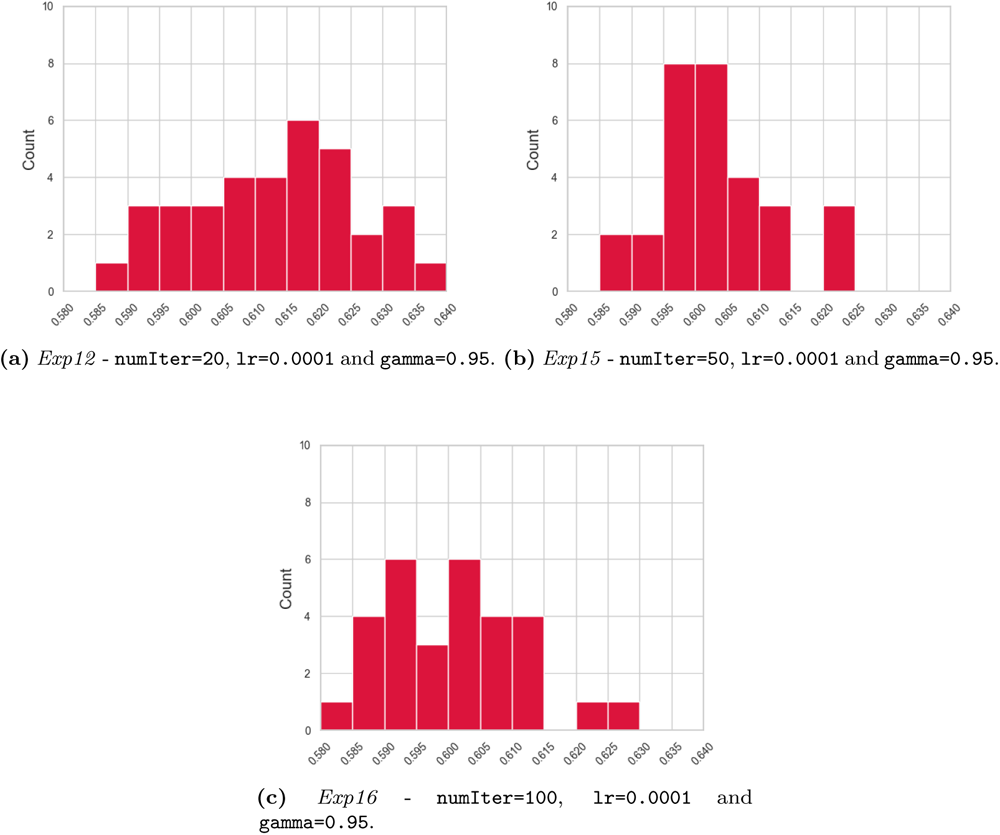
RL testing performances on Target 2 based on the numIter **hyperparameter values exploration.**. Histograms report distributions of the final number of cells obtained after each testing epoch. Each panel shows the performance of a trained model based on lr=0.0001, gamma=0.95, and different numIter values.

#### 4.4.4. Discussion on Target 2

As shown in subsubsection 4.3.4, the RL approach is capable for optimizing cell proliferation within the PalaCell2D simulation environment. These results confirm these results on Target 2, which contains the essential elements of a complex biological system, including both functional and structural aspects.

In summary, results on Target 2 show that the proposed RL approach is capable of:

- maximizing the final fraction of cells within the target at the end of a simulation;
- optimizing protocols of initial cell positions and compression stimuli coherently with the target and the modeling assumptions;
- expressing a range of training behaviors and performances with different combinations of learning and simulation hyperparameters (lr, gamma and numIter);
- learning and retaining knowledge within the trained ANN models, whose testing recapitulates training performance.

In subsubsection 4.4.1, the combination of lr and gamma values remarkably influences learning dynamics, leading two of the processes to very inferior performances than the rest and exhibiting a range of different balances between exploration and exploitation in general. In this experiment, performance is measured as *fraction_inside_*, the fraction of cells within the target area. The optimization controls not only the compression stimuli along each epoch but also the initial position of cells at the beginning of the simulation. This causes more dramatic oscillations in performance when the training explores divergent solutions. Indeed, the initial position of cells is poised to have a huge impact on the final performance (see Fig 10). Suppose the training sets a position that is too distant from the target. In that case, cells may have null overlap with it during that epoch, ultimately resulting in a null fitness value, even if compression stimuli are optimal. The numIter parameter, controlling the interaction frequency between the learning algorithm and the simulation environment, has a different impact than that shown in subsubsection 4.3.4. On Target 2, performances are lower when using higher numIter values (corresponding to fewer interactions between the learning process and the environment). Results suggest that, unlike in subsubsection 4.3.2, using numIter=50 and numIter=100 resulted in underfitting. Insufficient interaction with the environment reduced learning capacity, leading to insufficient exploration and adjustment of model parameters, resulting in poor performance [71]. Testing shows that the trained ANN models produce out-comes that are consistent with its training process (see subsubsection 4.4.3), confirming successful retainment of learnings acquired during training.

### 4.5. Comparison with GA-based optimized protocol generation

GAs are a class of computational methods inspired by natural evolution and selection principles, used to solve complex optimization and search problems [72]. They create a population of potential solutions, which evolves over successive generations through processes mirroring natural selection, crossover, and mutation. In each generation, individuals (solutions) are evaluated using a fitness function, and the fittest individuals are likelier to be chosen as parents for producing the next generation. GAs excel in solving problems with large, complex search spaces, where traditional optimization methods might falter, making them valuable in several fields, including the biomedical one [73]. This makes them perfect candidates for supporting a comparison of the proposed RL approach with state-of-the-art approaches. In addition, these two approaches stem from distinct problem-solving frameworks: RL from learning through interaction [74], and GA from evolutionary principles. Such a comparison benchmarks performance, highlighting which method converges more efficiently to a solution. It also contrasts their computational resource demands with RL’s sequential updates and GA’s parallel solution evaluations. To sustain the comparison, a GA-based pipeline has been built, resorting to the PyGAD library, an open-source Python library designed for the implementation of GAs [75]. GA runs were based on a population size of eight individuals, four parents mating, devising random mutation of solutions. Several open libraries for implementing the GA exist. The choice of PyGAD stems from its extreme flexibility in supporting user-defined fitness functions and complete control over GA evolution parameters. This facilitated the implementation of the interface between the optimization engine and the PalaCell2D simulator.

To provide a fair and broader comparison with RL performances, the GA runs were based on different numIter values, as they can be set for both methodologies. On the contrary, considering learning and evolution parameters would not sustain a consistent direct comparison. Indeed, parameters specific to the different optimization strategies are chosen after previous exploration (in the case of the RL-based approach) or following indications in the literature and documentation (in the case of the GA). Tab 5 reports the numIter values explored, and the last column reports the highest absolute fraction of cells inside the target across training for each experiment. The last column reports the RL (lr=0.0001 and gamma=0.95) performance with the same numIter value. The comparison considers GA-based and RL-based results based on different numIter values. Results show that the RL generates higher maximum fitness values for numIter=20 and numIter=50, while the GA surpasses it for numIter=100.

**Table 5:** Comparison between the highest fractions of cells obtained by GA and RL with different numIter hyperparameter values over Target 2. Each experiment corresponds to GA run with fixed population size (8) and number of parents mating (4) and the corresponding RL performance based on lr=0.0001, gamma=0.95 and different numIter values.

| Experiments | numIter | GA | RL |
| --- | --- | --- | --- |
| *Exp17* | 20 | 0.632 | 0.638 |
| *Exp18* | 50 | 0.629 | 0.635 |
| *Exp19* | 100 | 0.626 | 0.617 |

Fig 13a, Fig 13c and Fig 13e show the performance of the GA over 9 generations. The number of generations is chosen to support a fair comparison with the RL. The RL performance (Fig 13b, Fig 13d and Fig 13f) is evaluated on 70 epochs of training for each experiment, which corresponds to a total of 70 simulations of 2500 steps for Target 2. Each generation devises the simulation of 8 solutions for the GA. Thus, it takes 9 generations to complete a similar number of simulations (72) to the 70 simulations used by the RL trainings.

**Figure 13:**
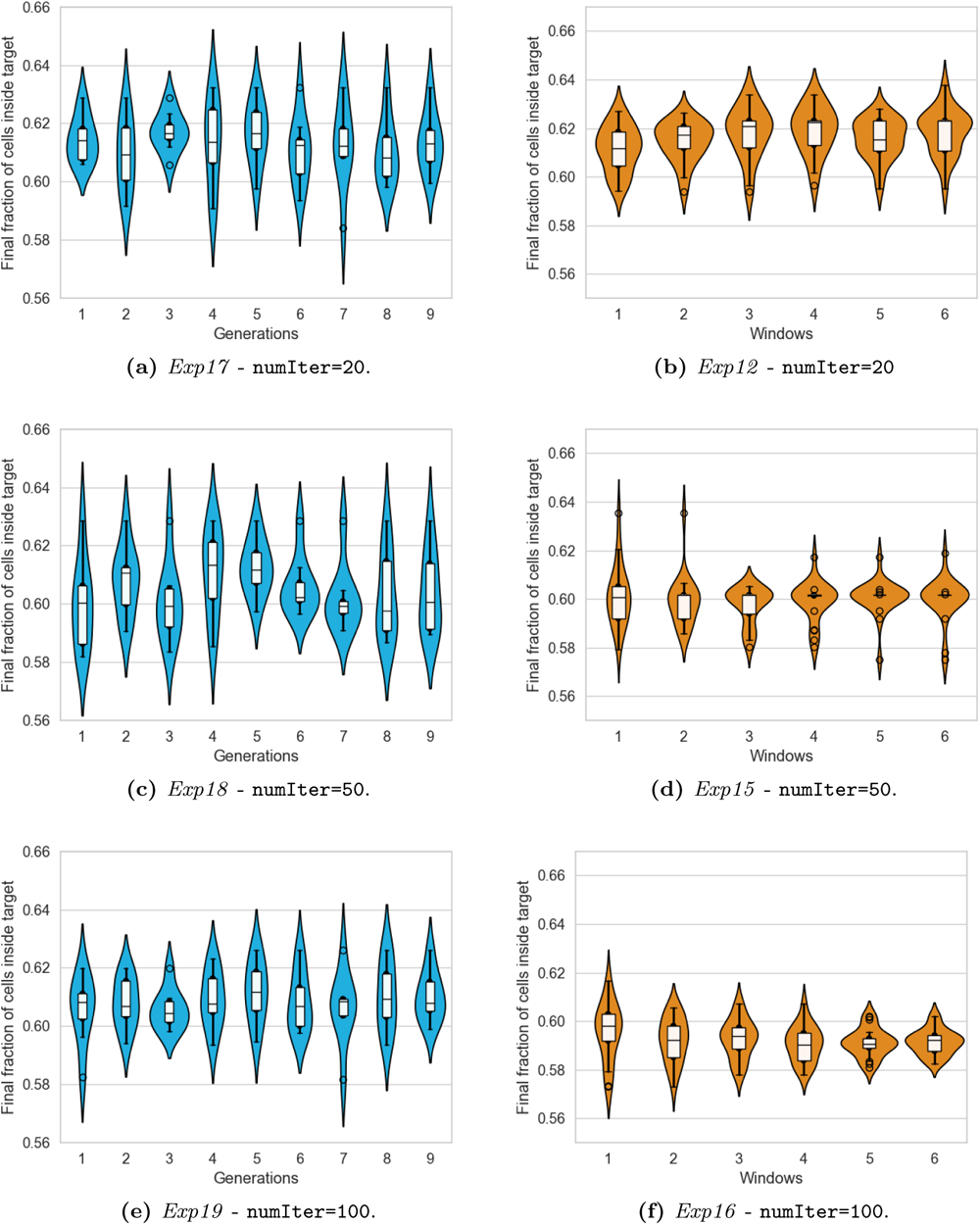
GA and RL performances on Target 2 - numIter **hyperparameter values exploration.** Each panel on the left shows the performance of a GA run with population size of 8, 4 parents mating and different numIter values over nine generations. Each panel on the right shows the performance of one training process based on lr=0.0001, gamma=0.95, and different numIter values over six 20-epochs windows.

Results show that the RL generates a higher *fraction_inside_* (0.638) than GA (0.632) for numIter=20 (see Tab 5). Fig 13 visualizes the corresponding evolutionary and learning processes: *Exp17* in Fig 13a and *Exp12* in Fig 13b, respectively, which both share the range of 0.60-0.62 for their median values across generations and windows. Yet, the RL training process generates a higher density of solutions in the upper part of the fitness values range, as indicated by the peaks of the density profiles, that gradually shift towards the upper boundary of the range (0.62). On the contrary, the GA generates solutions having a broader distribution of fitness values in all generations except for the third one, which exhibits a similar shape to the violin plots from the RL instead. This, combined with the steadily growing performance of the RL, shows that the capability to retain knowledge on the environment responses after stimulation allows the RL to dynamically adapt the generated solutions, converging to a higher-performance set of solutions. For numIter=50, the RL generates a higher *fraction_inside_*(0.635) than GA (0.629), but due to underfitting issues (see subsubsection 4.4.4) the training process stabilizes to lower performance levels (*Exp15* in Fig 13d), while the GA more extensively explores the solution space (*Exp18* in Fig 13c). Finally, for numIter=100, the RL generates a way lower *fraction_inside_* (0.617) than GA (0.626), and overall poorer performance across training (*Exp16* in Fig 13f), while the GA keeps exploring the space effectively (*Exp19* in Fig 13e).

### 4.6. Computational comparison

Eventually, this section aims to compare the computational complexity of all experimental setups briefly. All experiments ran on an AMD Ryzen 9 5950X 16-Core 2.2GHz processor with 64GB RAM. The execution environment leveraged the container platform Singularity [76].

Table 6 summarizes CPU times for relevant tasks across optimization targets and experiments, distinguishing results for RL and GA.

**Table 6:** Typical CPU times of RL epochs and GA generations for Target 1 and Target 2.

| Approach | Target | Task | Typical CPU time (s) |
| --- | --- | --- | --- |
| RL | 1 | Epoch | ~60 |
| RL | 2 | Epoch | ~400 |
| GA | 2 | Generation | ~200 |

Results display how the RL pipeline requires more computational time than the GA one. It is worth noting that the extra cost should be compensated not only by better results, as discussed in Section 4.5, but also by the potential re-usage of the learned protocol and trained ANN model.

## 5. Conclusions

In our study, we developed a computational DSE method to optimize the biofabrication of epithelial sheets, improving upon previous research [18]. This method utilizes a variant of the A2C RL algorithm alongside the PalaCell2D simulator [20]. We validated it through two main objectives: increasing total cell count and enhancing cell density in specific areas. The RL approach, influenced by compression and initial cell placement, showed varied learning dynamics and performance in both training and testing phases.

The interplay of hyperparameters like lr, gamma, and numIter is crucial for balancing quick adaptation and thorough exploration. Notably, numIter’s impact changes depending on the objective, highlighting the need for cautious evaluation to avoid overfitting or underfitting.

In comparison to a GA method for one of the targets, the RL approach exhibited better performance when under optimal hyperparameter setting. This underscores the importance of meta-optimization in model-based OvS of complex simulated environments [17]. However, the RL’s capability for learning distinguishes it from evolutionary optimization strategies in general, and makes it particularly apt for real-world scenarios. In fact, as it integrates simulation with optimization and RL training, this approach couples compuational DSE with learning, supporting subsequent generalization to different applications.

This study, while limited to a single A2C variant and ANN architecture, establishes a foundation for optimizing *in silico* culture processes. Future research should focus on creating a versatile framework accommodating different algorithms, simulators, and use cases to increase computational efficiency and widen its use in computational biology. The reusability of both the optimized protocol and the trained ANN, along with their demonstrated optimization and learning capabilities, supports the usage and further development of RL strategies for the advancement of TE and RM biofabrication.

## 6. Data Availability Statement

The source code and data used to produce the results and analyses presented in this manuscript are available from the public GitHub repository https://github.com/smilies-polito/rLotos.

## 7. Acknowledgements

We acknowledge the precious collaboration and support provided by PalaCell2D team, Dr. Raphaël Conradin, and Prof. Bastien Chopard from the Scientific and Parallel Computing (SPC) Group at the University of Geneva, Geneva, Switzerland.

## Appendices

### Appendix A. Acronyms

**A2C** Synchronous Advantage Actor-Critic

**ABM** Agent-Based Model

**AI** Artificial Intelligence

**ANN** Artificial Neural Network

**CFD** Computational Fluid Dynamics

**DoE** Design of Experiment

**DL** Deep Learning

**DRL** Deep Reinforcement Learning

**DSE** Design Space Exploration

**FEM** Finite Elements Model

**GA** Genetic Algorithm

**ML** Machine Learning

**MLP** Multi-Layer Perceptron

**MSE** Mean Squared Error

**OFAT** One Factor At a Time

**ODE** Ordinary Differential Equation

**OvS** Optimization via Simulation

**PN** Petri Nets

**RL** Reinforcement Learning

**RM** Regenerative Medicine

**TE** Tissue Engineering

### Appendix B. Framework setup parameters

This section overviews relevant parameters to the framework setup, including simulation and learning environment configuration and the management of output files for visualization and performance metrics calculations.

**Simulation setup** The PalaCell2D simulation configuration file holds different types of simulation parameters. Some parameters control the simulation duration:

- numIter controls the number of simulation steps (or iterations) to perform.
- stopAt indicates the simulation step (or iteration) when to stop the simulation.

Another set of parameters is the ones controlling the simulation evolution. The biofabrication protocol sets their values according to the actions generated by the RL algorithm along the training. This work includes a new parameter to the ones provided in PalaCell2D: initialPos, the position of the first cell generated in the simulation space.

- initialPos sets the position of the initial cell, indicating its x and y coordinate values.
- compressionAxis sets the axis for external force application, with either ’X’ or ’Y’ as values.
- comprForce indicates the intensity of the external force applied.

Another set of parameters in the configuration supports the management of input and output files. Both types are .vtp files from the Visualization Toolkit VTK [77].

- initialVTK indicates the name of the input .vtp file from which to restore the simulation (if this value is missing, the simulation starts from the beginning).
- finalVTK indicates the name of the generated output .vtp file.

**Learning environment setup** This work implements the simulator interface (see section 3) to use PalaCell2D simulations as a learning environment. Moreover, it relies on the configure method, called by the implemented environment, to create the configuration file needed by the PalaCell2D simulator. The implemented PalaCell2D environment has the following parameters to interface the learning process.

- width and height: the width and height in pixels of the observations provided to the learning model, with a default value of 300.
- numIter: the number of simulation steps per learning epoch, with a default value of 20.
- max_iterations: the maximum number of total simulation steps, with a default value of 4200.

**Output management** Performance evaluation relies on computing defined metrics over the generated output. Data saving occurs every five training epochs. The helper file vtkInterface.py contains the following methods to read PalaCell2D outputs.

- read_cell_num: reads the number of cells in the simulation space from the simulation output file.
- create_png_from_vtk: reads the simulation output file and produces a visualization of the simulation space. This visualization centers on the cellular aggregate and not on the simulation space.
- create_decentered_pil_image: reads the simulation output file and produces a visualization of the simulation space. This visualization centers on the simulation space.
- add_target: adds a circular target with given center coordinates and radius to a chosen image.
- count_target_points: reads the simulation output file and returns the number of cellular vertices inside and outside a specified target.

### Appendix C. Simulator interface

The simulator interface contains the set of required functions and a format for data exchange to fulfill the specific objectives and requirements of different applications. Each specific application requires an implementation of the simulator interface, where the necessary functions are implemented to interact with the chosen simulator. Each implementation of the simulator interface creates an environment based on the *environment* file, whose structure follows the one proposed in [78]. The *environment* file sets the ANN model structure (see subsection 3.2) and the training process parameters for the target application.

The simulator interface includes the following required functions to advance the training process.

- reset: prepares the environment for the next episode starting from an initial state;
- render: returns an image showing the environment state;
- adapt_action: transforms the actions provided by the ANN in a format that can be used by the step method;
- step: acts on the environment with the generated actions and provides an observation of the environment, the reward value, and a boolean value that tells whether the environment has reached a terminal state or not to the training process.

The simulator interface also includes the following set of required functions to manage data produced by the training process and keep track of performance.

- save_performance: indicates whether or not to collect performance indexes related to the environment into environment variables.
- get_performance: saves the performance indexes in a checkpoint file.
- load_performance: restores the performance indexes in the respective environment variables when restoring the training from a previous state through checkpoint files.
- check_performance: indicates whether or not to save the collected performance indexes in a checkpoint file.
- data_to_save: indicates which data to save in a checkpoint file that is accessible after the training.
- load_data_to_save: restores the collected data in the respective variables inside the environment when restoring the training from a previous state through checkpoint files.

### Appendix D. Nested training process

To tackle Target 2 (see subsection 4.4), a helper training class allows synchronizing the two nested subprocesses. This class has the same structure as the one already presented in subsection 3.4, with the addition of a pipe allowing communication between the two subprocesses. The two environments perform the steps described in Algorithm 1 during the training process.

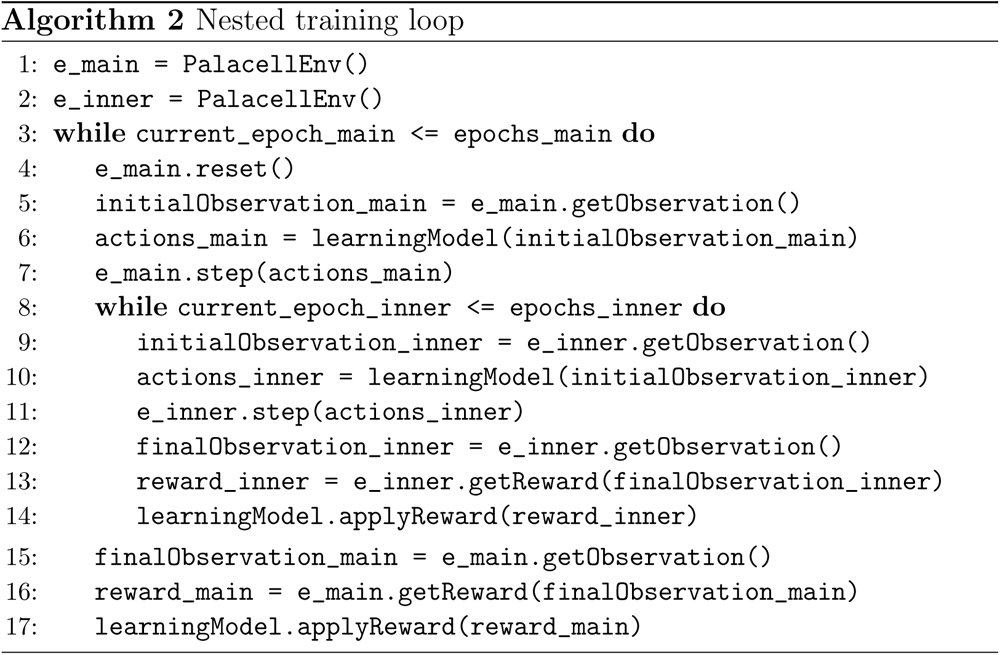

This flow leverages communication mechanisms between subprocesses, which supports the steps illustrated in Alg. 2. After instantiation of the main and inner learning environments (Alg. 2, lines 1-2), the training creates a simulation configuration file for the inner training with the same values as in the previous experiment (see subsection 4.3). The initialPos parameter it is not constant but rather set by actions generated by the learning model. The main subprocess runs its training process (Alg. 2, lines 3-7) and sends the simulation configuration to the second training process through the pipe. The second subprocess uses it as the starting configuration file, then runs its training process (Alg. 2, lines 8-14). This training process runs the same number of simulation iterations per epoch as subsection 4.3 (numIter=20) for a total of 2500 simulation iterations. When the second training process terminates, its subprocess sends the generated output to the main subprocess through the pipe. The main subprocess reads the output, computes performance metrics, and produces an observation for the main training process (Alg. 2, line 15). The main training process learns through observation and the corresponding reward (Alg. 2, lines 15-17). Then, it proceeds to the next epoch.

